# EPHA1 and EPHB4 tyrosine kinase receptors regulate epithelial morphogenesis

**DOI:** 10.1101/2024.07.15.603563

**Authors:** Noémie Lavoie, Anaëlle Scribe, François J.M. Chartier, Karim Ghani, Alexandra Jetté, Sara L. Banerjee, Manuel Caruso, Mélanie Laurin, Andrew Freywald, Sabine Elowe, Patrick Laprise, Nicolas Bisson

**Affiliations:** Centre de recherche du Centre Hospitalier Universitaire (CHU) de Québec-Université Laval, Division Oncologie, Québec, QC, Canada; Centre de recherche sur le cancer de l’Université Laval, Québec, QC, Canada; PROTEO-Quebec Network for Research on Protein Function, Engineering, and Applications; Institut de recherches cliniques de Montréal, Montréal, QC, Canada; Department of Molecular Biology, Medical Biochemistry and Pathology, Université Laval, Québec, QC, Canada; Department of Pathology and Laboratory Medicine, College of Medicine, University of Saskatchewan, Saskatoon, SK, Canada; Department of Pediatrics, Université Laval, Québec, QC, Canada

## Abstract

Organ formation and homeostasis require the coordination of cell-cell adhesion, epithelial cell polarity and orientation of cell division to organize epithelial tissue architecture. We have previously identified proximity protein networks acting downstream of members of the EPH family of tyrosine kinase receptors and found within these networks an enrichment of components associated with cell morphogenesis and cell-cell junctions. Here, we show that two EPH receptors, EPHA1 and EPHB4, are localized to the basolateral domain of Caco-2 cells in spheroidal cultures. Depletion of either EPHA1 or EPHB4 disrupts spheroid morphogenesis, without affecting cell polarity, but via randomizing mitotic spindle orientation during cell division. Strikingly, EPHA1 and EPHB4 exert this function independently of their catalytic activity but still requiring EFN ligand binding. Consistent with this, the most abundantly expressed EPHB4 ligand in Caco-2 cells, EFNB2, is also compartmentalized at the basolateral domain in spheroids, and is required for epithelial morphogenesis. Taken together, our data reveal a new role for EPHRs in epithelial morphogenesis.

## INTRODUCTION

Epithelial tissues act as barriers with selective permeability, creating in organs compartments with distinct biochemical compositions. This primary function of epithelial tissues relies on the *zonula adherens*, which provides cell-cell cohesion, as well as tight junctions that seal the intercellular space and preclude passive diffusion across epithelial sheets (Díaz-Díaz *et al*, 2020). In addition, epithelial tissues fulfill their role by guiding vectorial transport and by performing spatially oriented secretion of selected molecules. The unidirectional nature of these functions requires the asymmetric distribution of many cell components. This structural organization is referred to as epithelial cell polarity, which is characterized by the presence of an apical domain facing the external environment (or a lumen), a lateral domain contacting neighbouring cells, and a basal domain anchored to the basement membrane – a specialized layer of the extracellular matrix (Buckley & St Johnston, 2022; Martin *et al*, 2021). Epithelial tissue morphogenesis generates functional structures sustaining organ physiology such as tubules in kidneys, alveoli in lungs, acini in mammary glands, and villi in the small intestine (Schöck & Perrimon, 2002; Bernascone *et al*, 2017). Acquisition and maintenance of these 3D units depend on the fine coordination between cell-cell adhesion, epithelial cell polarity, and the orientation of cell division. In simple epithelia, the axis of cell division is perpendicular to the plane of the tissue, thereby ensuring that both daughter cells are positioned side-by-side within monolayered epithelial sheets (Bergstralh *et al*, 2017).

EPH receptors (EPHRs) constitute the largest family of receptor tyrosine kinases. They are divided into two groups: EPHAs (A1–8, A10) and EPHBs (B1–4, B6), mainly based on their preferences for binding membrane-bound ephrin (EFN) ligands of the A (EFNA1–5) or B (EFNB1–3) subgroups. EFNAs are tethered to the cell membrane by a glycosylphosphatidylinositol (GPI) anchor, whereas EFNBs are single-pass transmembrane proteins with a short cytoplasmic tail (Kania & Klein, 2016). Consistent with the membrane localization of EFNs, EFN-EPHR interactions are mainly involved in short-range cell-to-cell communications, requiring direct contacts and leading to bidirectional signaling events mediated by both EPHRs and EFNs (Cayuso *et al*, 2015). Many reports have demonstrated that both EPHR-mediated forward signaling and EFN-mediated reverse signals are required to accomplish a given function (Boyd *et al*, 2014).

EPHR signaling is mediated via an intracellular region composed of four highly conserved domains that enable interactions with cytoplasmic effectors: a juxtamembrane segment containing two key tyrosine (Tyr) residues, a tyrosine kinase domain, a sterile-α motif (SAM) domain, and a PSD-95, DLG1, ZO-1 (PDZ) domain-binding motif (PBM). Most of the interactions described to date require activation of the EPHR Tyr kinase function and its phosphorylation on cytoplasmic Tyr residues (Oughtred *et al*, 2021), highlighting the importance of phospho-Tyr (pTyr)-mediated signaling for EPHR-dependent cellular functions. EFN-EPHR signaling modules use complex signaling networks to control a vast array of biological processes impacting the formation and maintenance of tissue organization (Arcas *et al*, 2020). For instance, EFN-EPHR signaling is required to define cell boundaries within and between different tissues by regulating cell adhesion and repulsion (Kania & Klein, 2016). In addition, EPHR-dependent regulation of cell shape, adhesion and migration is crucial for maintaining plasticity and regenerative capacity (Klein, 2004). The sole EFN-EPHR pair in *D. melanogaster* signals to restrict proliferation and to control spindle orientation and the plane of division in symmetrically dividing neuroepithelial cells (Franco & Carmena, 2019a). It remains to be determined whether EFN-EPH signaling plays a similar role in maintaining epithelial tissue organization and homeostasis in vertebrates. Supporting this possibility, we previously defined the proximity networks of selected EPHRs and found several proteins controlling cell-cell adhesion and epithelial cell polarity (Banerjee *et al*, 2022), which can both determine spindle orientation (Wang *et al*, 2018; Tuncay & Ebnet, 2016). Further investigating this question is of high significance given that dysregulation of EPH-EFN signaling, spindle orientation, or cell polarity contributes to cancers of epithelial origin (Anderton *et al*, 2021).

EFN ligands and EPH receptors are expressed in several epithelial tissues, including the intestinal epithelium where they contribute to the appropriate positioning of intestinal epithelial cells along the crypt-villus axis (Cortina *et al*, 2007; Clevers & Batlle, 2006; Batlle *et al*, 2002). The adenocarcinoma Caco-2 cell line is useful to study several aspects of intestinal epithelial cells biology and to model the morphogenesis of simple epithelia. In 2D culture systems, these cells are proliferative and unpolarized at sub-confluency. Upon reaching confluence, Caco-2 cells exit the cell cycle and initiate a differentiation program that extends over several days and results in the formation of adherens and tight junctions, polarization along the apical-basal axis, and expression of intestine-specific proteins. When seeded in a 3D Matrigel matrix, Caco-2 cells form spheroids made of a monolayer of polarized cells with their apical surface facing a single central lumen (Jaffe *et al*, 2008). Caco-2 cells express several EPHRs with *EPHA2*, *-A4*, *-B4* and *-A1* being the most abundantly expressed (Zecha *et al*, 2020; Uhlén *et al*, 2015). However, the subcellular distribution and function of these receptors in these cells remain unexplored. In the present study, we used Caco-2 in 3D culture to assess the role of selected EPHRs in spheroid formation and organization, thus furthering our understanding of the role of these receptors in epithelial tissue morphogenesis.

## RESULTS

### EPHA1 and EPHB4 are expressed basolaterally in Caco-2 spheroids

RNA expression profiling revealed that Caco-2 cells mainly express EPHA2, EPHA4, EPHB4 and EPHA1 (Figure 1A). To explore the role of EPHR signaling in Caco-2 cell spheroid formation, we first monitored expression of endogenous EPHA2, -A4, -B4 and -A1 receptors in sub-confluent proliferating cells, cell cycle-arrested confluent monolayers, and post-confluent differentiating Caco-2 cells grown in 2D. All four receptors were detected in sub-confluent Caco-2 cell cultures (Figure 1B). However, levels of EPHA2 and EPHA4 decreased at confluency and was further reduced in 5 days post-confluent monolayers. This suggests that EPHA2 and EPHA4 are mainly required in proliferating cells and that Caco-2 cells likely differentiate and polarize without a major contribution of these receptors. Hence, we focused our analysis on EPHA1 and EPHB4, which displayed a sustained expression in confluent and post-confluent Caco-2 cells. EPHA1 and EPHB4 were both localized at the lateral and basal membranes in polarized Caco-2 cells grown in 3D and forming hollow spheroids (Figure 1C). These receptors were excluded from tight junctions, which are marked by ZO-1 staining and delimit the boundary between the lateral and apical domains. Moreover, EPHA1 and EPHB4 did not overlap with EZRIN (EZR), which decorates the lumen-lining apical free surface. Overall, these data show that EPHA1 and EPHB4 are expressed in differentiated Caco-2 cells and display a polarized distribution in these cells by specifically localizing to the basal and lateral domains.

**Figure 1.**
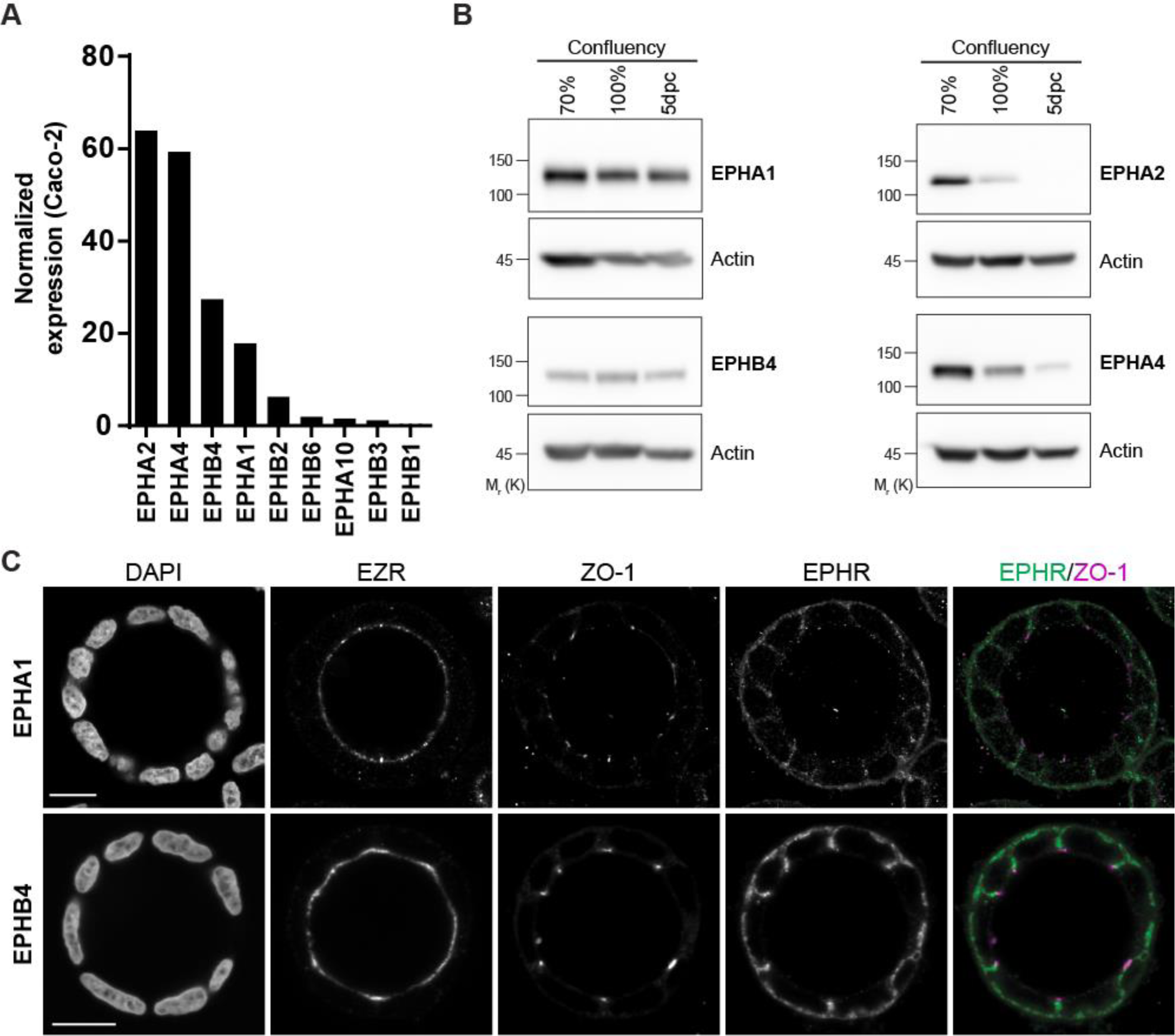
EPHA1 and EPHB4 are expressed basolaterally in Caco-2 spheroids. (A) RNA expression levels of all human EPHRs in Caco-2 cells were extracted from the Human Protein Atlas database and displayed in a decreasing order. (B) Western blot analysis of endogenous EPHA1, EPHA2, EPHA4 and EPHB4 in sub-confluent (70%), confluent (100%) and post-confluent (5dpc; 5 days post-confluency) Caco-2 cells cultured as monolayers. Representative images of 3 experiments are shown. (C) Confocal images displaying a representative example of EPHA1 and EPHB4 expression in 3D Caco-2 spheroid cysts. Single cells were seeded on Matrigel and grown for 6 days. Expression of apical (EZR) and tight junction (ZO-1) markers is shown, along with nuclear DAPI staining (scale bar: 15 µm).

### EPHA1 and EPHB4 are required for Caco-2 cell spheroid morphogenesis

To test whether EPHA1 and EPHB4 are required for morphogenesis of Caco-2 cell spheroids, we individually depleted their expression using two independent shRNAs prior to cell seeding in Matrigel. We found that EPHA1 or EPHB4 depletion led to the formation of disorganized Caco-2 cell cysts with multiple atrophic lumens in over 70% of cases, as shown by EZR distribution (Figure 2A-B). This phenotype is reminiscent to what is observed following depletion of several epithelial cell polarity regulators, including the atypical kinase PKCι (PRKCI) in Madin-Darby Canine Kidney (MDCK) epithelial cell cysts (Linch et al, 2014) (Durgan *et al*, 2011). Similarly, we found that knockdown of PKCι in Caco-2 cells led to the production of aberrant multi-lumen structures (Figure 2A-B). Knockdown efficiency was confirmed by Western blotting for each shRNA sequence (Figure 2C). Our observations raised the possibility that EPHA1 and/or EPHB4 support the formation of organized Caco-2 cell spheroids by contributing to epithelial cell polarization. We thus examined the distribution of apical (PARD6, EZR), tight junction (ZO-1), and basolateral (SCRIB, E-CDH) markers in control and EPHA1- or EPHB4-depleted spheroids. We observed proper segregation of most of these apical, tight junction, and lateral proteins in absence of EPHA1 and EPHB4, showing that individual Caco-2 cells are polarized despite the fact that they produce disorganized structures (Figure 3, Figure S1). However, we observed a mild EZR delocalization to the basal region in 64.3% of EPHA1-depleted spheroids, while EPHB4 depletion had little to no effect (Figure S2). This suggests that EPHA1 plays a role in restricting EZR to the apical membrane, but is not required to establish the apical-basal axis. Collectively, these results show a role for EPHA1 and EPHB4 in the formation of Caco-2 cell spheroids, a model for morphogenesis of single-layered epithelial, without having a major impact on epithelial cell polarity.

**Figure 2.**
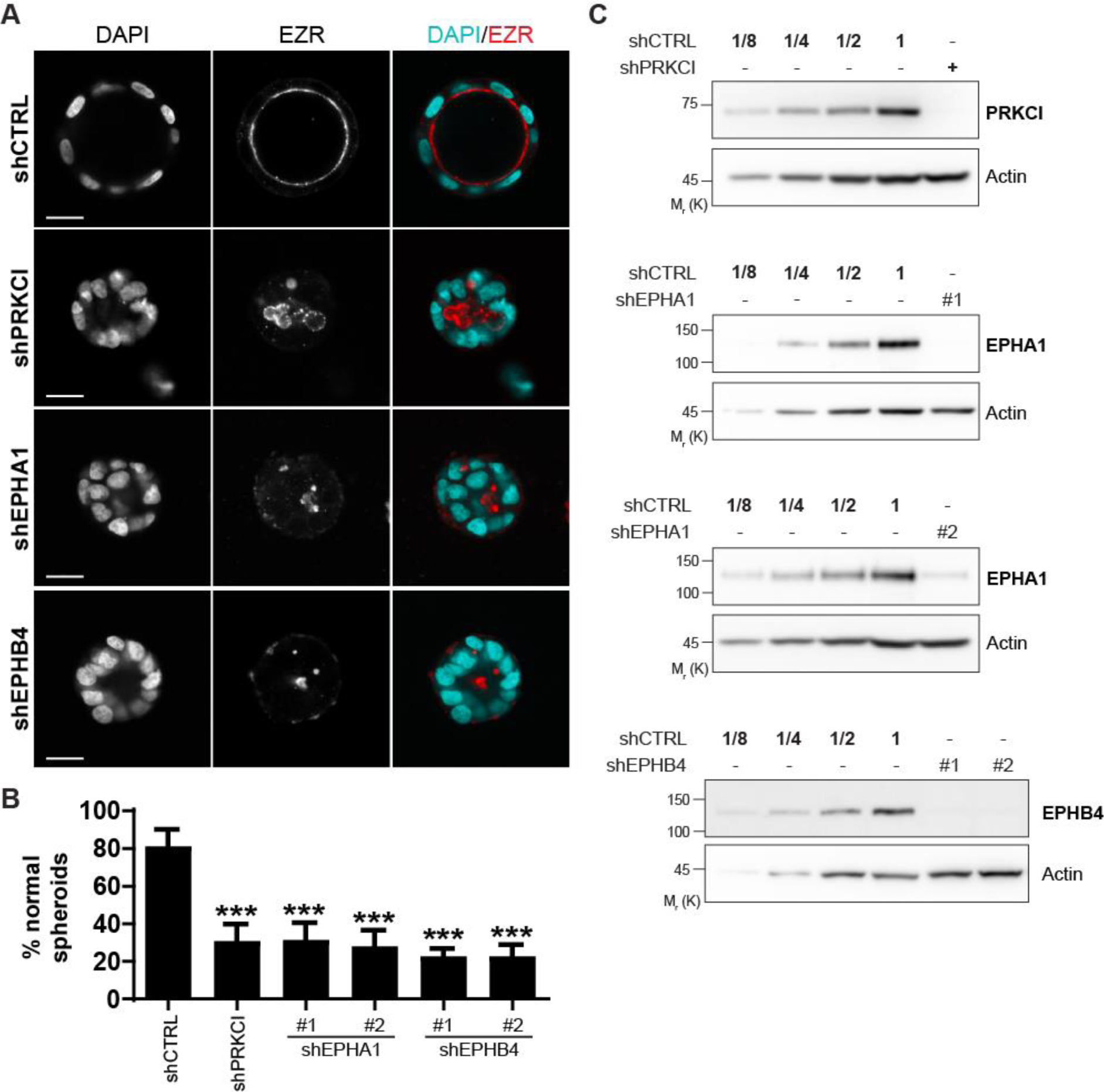
EPHA1 or EPHB4 depletion leads to defects in Caco-2 spheroid formation. (A) Representative images of Caco-2 spheroids morphology following depletion of PKCι, EPHA1 or EPHB4. Two independent shRNAs were used against EPHRs. EZR apical marker highlights the lumen border, along with nuclear DAPI staining. Single cells were seeded on Matrigel and grown for 6 days (scale bar: 15 µm). (B) Quantification of the percentage of Caco-2 spheroids displaying a wild-type morphology (i.e. single, round lumen judged by EZR staining, surrounded by a monolayer of cells) following depletion of PKCι, EPHA1 or EPHB4. The mean of three independent experimental replicates is shown. Error bars indicate SD (***p≤0,001; ANOVA with Tukey’s correction for multiple comparison; n=400 for shCTRL and shPKCι, and n=150 for each shEPH). (C) Western blot analysis of endogenous PKCι, EPHA1 or EPHB4 following shRNA depletion. Successive dilutions of control are indicated to estimate depletion efficiency, where 1 is equivalent to 50 µg. Representative images of 3 experiments are shown.

**Figure 3.**
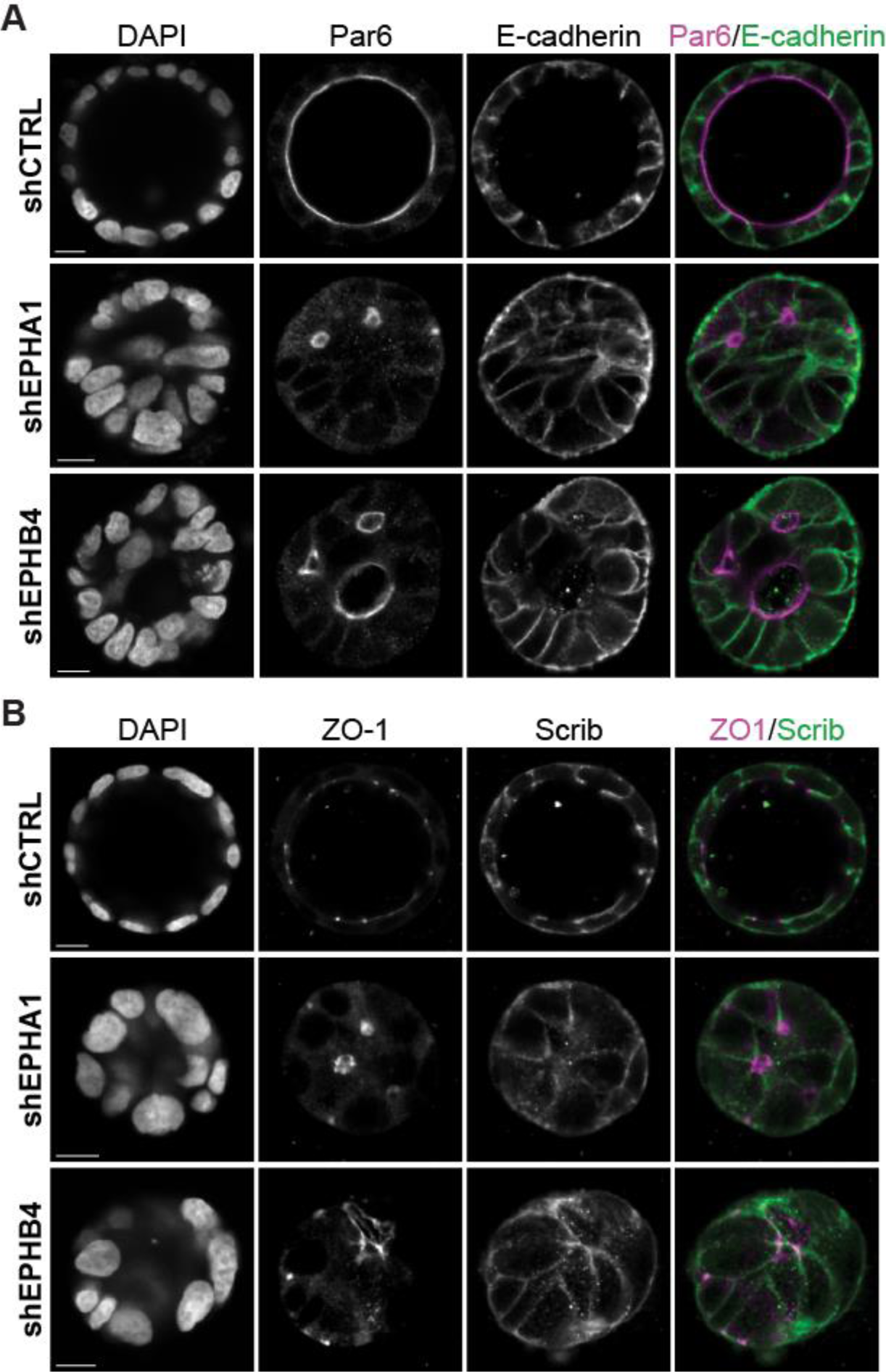
EPHA1 and EPHB4 depletion does not affect apical and basal polarity markers localization. (A-B) Representative images showing polarity proteins subcellular localization in Caco-2 spheroids following depletion of EPHA1 and EPHB4. Single cells were seeded on Matrigel and grown for 6 days. PARD6 (Par6) apical marker highlights the lumen border, ZO-1 is located at tight junctions and E-CDH and SCRIB highlight basolateral membranes, along with nuclear DAPI staining (scale bar: 12 µm). Corresponding Western blots shown in Figure S1A.

### EPHA1 and EPHB4 depletion leads to mitotic spindle orientation defects

In monolayered epithelia, the axes of cell division are oriented perpendicular to the plane of the tissue so that both daughter cells reside side-by-side within the epithelial sheet instead of piling up on top of one another. This contributes to the maintenance of the architecture of these tissues (Bergstralh *et al*, 2017). Similarly, Caco-2 cells divide at an angle of 90° relative to the center of the spheroid in order to establish a monolayered epithelial structure (Jaffe *et al*, 2008). Randomization of spindle pole orientation in Caco-2 cells results in the formation of disorganized cysts containing numerous lumens (Rodriguez-Fraticelli *et al*, 2010a; Hao *et al*, 2010a). This raises the possibility that EPHA1 and EPHB4 control Caco-2 spheroid morphogenesis by regulating mitotic spindle orientation. To explore this possibility, we measured angles between centrosome-attached spindles using alpha-tubulin (TUBA1A) staining and the center point of spheroids. We found that EPHA1- or EPHB4-depleted cells divide at random angles (average of 47°, median of 55° for shEPHA1 and average/median of 45° for shEPHB4), whereas control cells divide at an average angle of 73° (median 79°), much closer to the expected theoretical 90° (Figure 4A-C, Figure S1). We also observed that EPHRs are maintained at the basolateral membrane in dividing control cells (Figure 4D), suggesting that these receptors could contribute to anchoring the spindle to the cell cortex. These data provide evidence that spindle orientation defects cause inadequate cellular organization in EPHR-depleted Caco-2 spheroids.

**Figure 4.**
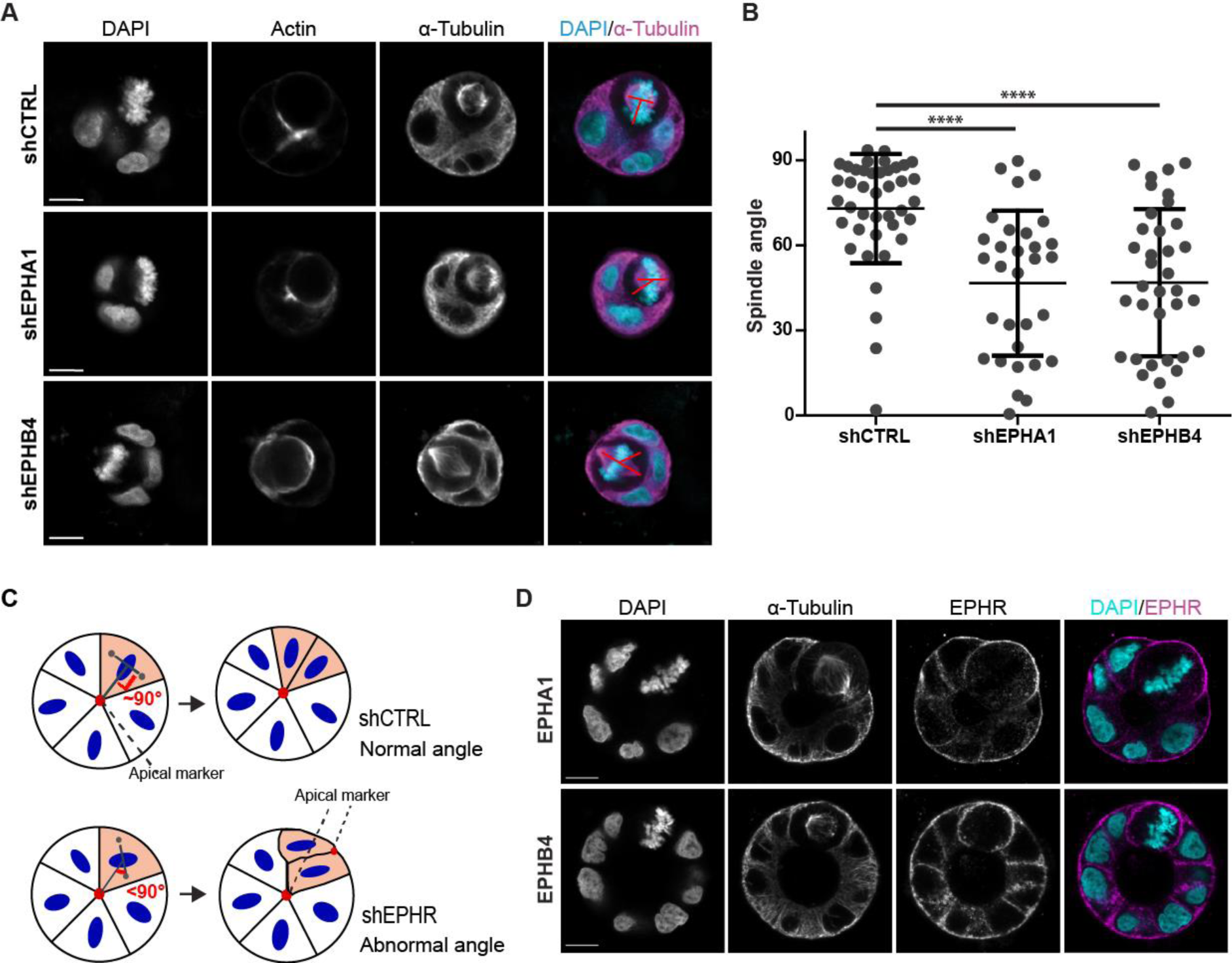
EPHA1 and EPHB4 depletion leads to mitotic spindle orientation defects. (A) Representative images showing orientation of dividing cells in Caco-2 spheroids following depletion of EPHA1 and EPHB4. Single cells were seeded on Matrigel and grown for 2 days. Actin highlights the apical patch, and α-tubulin shows microtubules, along with nuclear DAPI staining. (scale bar: 10 µm) (B) Graph showing the mean angle of mitotic spindle in dividing cells. Angle was measured between a first line (joining the centrosomes) and a second line (joining the center of the cyst and the first line). Error bars indicate SD (***p<0,0005; Kruskal-Wallis statistic test; n=41 for shCTRL n=31 for shEPHA1 and n=36 for shEPHB4). (C) Schematic representation of the impact of improper cell division orientation on spheroid formation. (D) Confocal images displaying a representative example of EPHA1 and EPHB4 expression in dividing cells in 3D Caco-2 cysts. Single cells were seeded on Matrigel and grown for 6 days. Staining of ɑ-tubulin highlight dividing cell, along with nuclear DAPI staining (scale bar: 10 µm). Corresponding Western blots shown in Figure S1B.

### EPHA1- and EPHB4-regulated spheroid morphogenesis requires an intact ligand binding domain, but is independent from their catalytic activity

To confirm the specificity of phenotypes seen with loss-of-function experiments and to better understand how EPHA1 and EPHB4 each act in coordinating epithelial morphogenesis, we performed rescue experiments. We used retroviral infection to express shRNA-resistant EPHA1-GFP and EPHB4-GFP at near endogenous levels in EPHA1- and EPHB4-depleted cells, prior to seeding in Matrigel. EPHA1-GFP or EPHB4-GFP were properly localized at lateral and basal membranes, and their expression in presence of endogenous receptors did not affect spheroid formation (Figure S3). Importantly, expression of a cognate shRNA-resistant GFP-tagged WT EPHR in EPHA1 or EPHB4 knockdown cells significantly restored the formation of normal Caco-2 spheroids with a monolayer of cells surrounding a single lumen (p<0.0001; ANOVA with Dunnett’s correction for multiple comparison) (Figure 5). This confirms that the phenotype described (Figure 2) is dependent on EPHR expression. Likewise, kinase inactive EPHA1(K656A)-GFP and EPHB4(K647A)-GFP were also able to replace endogenous receptors and restore the ability to form hollow spheres composed of a monolayered epithelial sheet (p<0.001) (Figure 5), indicating that EPHRs’ catalytic activity is not required for the process.

**Figure 5.**
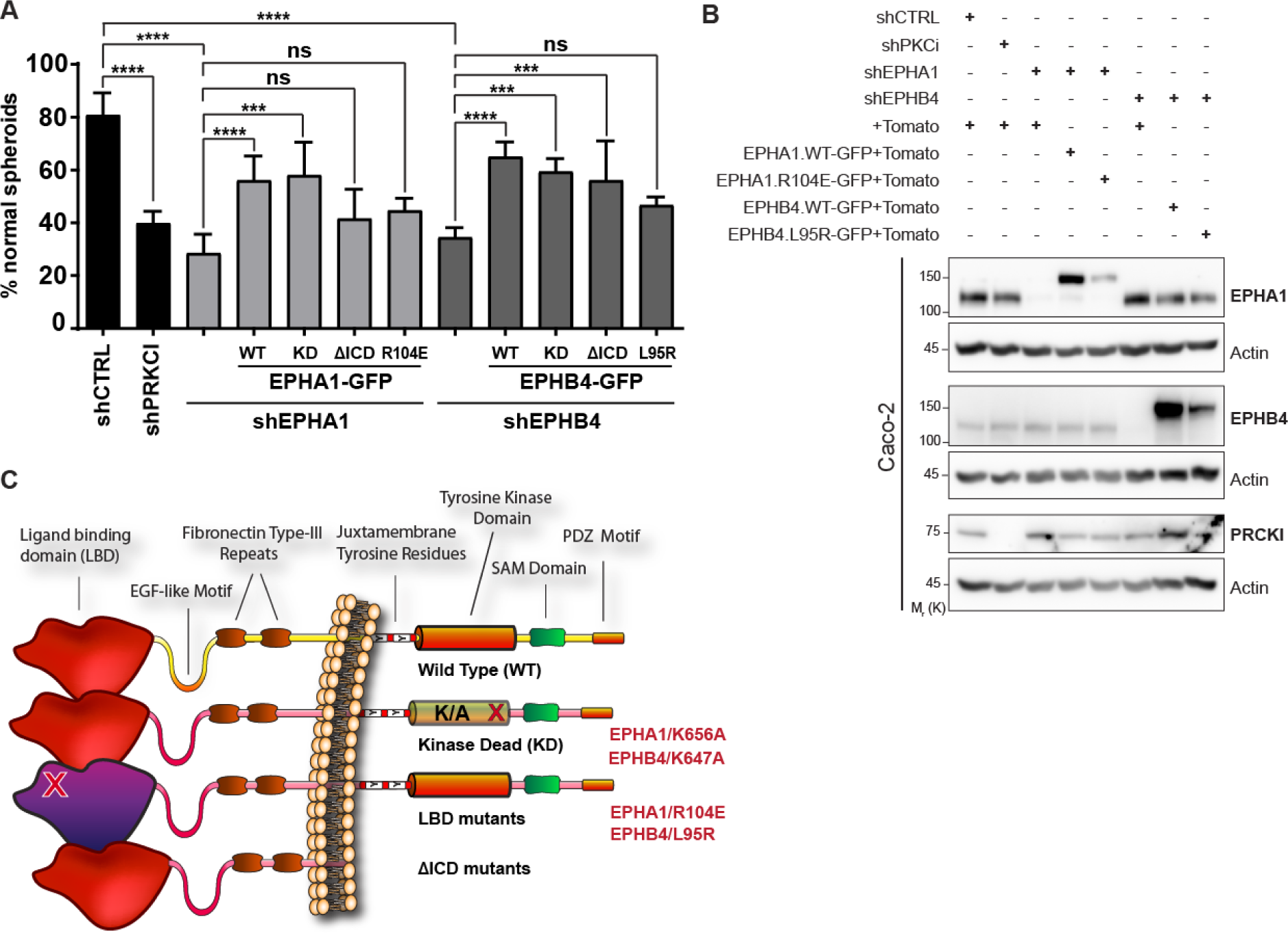
EPHA1- and EPHB4-regulated spheroid morphogenesis requires an intact ephrin ligand binding domain, but is independent from their catalytic activity. (A) Quantification of the percentage of Caco-2 spheroids displaying a wild-type morphology (i.e. single, round lumen judged by EZR staining, surrounded by a monolayer of cells) following rescue with EPHA1 or EPHB4 WT or mutant forms. The mean of at least three independent experimental replicates is shown. Error bars indicate SD (*p≤0,05, **p≤0,01, ***p≤0,001, ****p≤0,0001; ANOVA with Dunnett’s correction for multiple comparison; n=550 for shCTRL+Tomato and shPKCι+Tomato, n=450 for shEPHA1+Tomato, shEPHB4+Tomato and EPHR-WT-GFP rescue, n=250 for EPHR-ΔICD-GFP rescue and n=150 for EPHR-KD-GFP, EPHA1-R104E and EPHB4-L95R rescue) (B) Western blot analysis of EPHA1 and EPHB4 endogenous levels in knockdown and rescue experiments in Caco-2 cells cultured as monolayers. Representative images of at least 3 experiments are shown. (C) Schematic representation of EPHR mutants.

To assess the possible requirement of EPHA1/B4 functional domains, we pursued rescue experiments using EPHR mutants defective in either their ephrin ligand binding domain (LBD) (Singh *et al*, 2018; Himanen *et al*, 2009; Chrencik *et al*, 2006), or truncated to eliminate their entire intracellular domain (ΔICD), which includes tyrosine kinase and SAM domains, as well as a PDZ binding motif. We found that LBD-defective EPHA1(R104E)-GFP and EPHB4(L95R)-GFP were unable to restore spheroid morphology in EPHA1- and EPHB4-depleted cells (Figure 5). We also discovered that only EPHA1 requires its intracellular domain, as EPHB4(ΔICD)-GFP was capable of restoring spheroid morphology in EPHB4-depleted cells (p<0.001) (Figure 5). Together, these results suggest that ephrin binding to these receptors is required for the establishment or maintenance of spheroid morphology. They also indicate that, while EPHA1/B4 tyrosine kinase activity is not involved in the process, the two receptors differ in their mode of action, with only EPHA1 requiring its intracellular functional domains. Thus, signaling initiated by EFNs in their host cells is likely to act as the major mediator of EPHB4 function.

### EFNB2 is required for Caco-2 spheroid morphogenesis

According to The Human Protein Atlas database, EFNB2, EFNA1 and EFNA4 are the most abundantly expressed EFN ligands in Caco-2 cells (Uhlén *et al*, 2015) (Figure 6A). As only EFNBs were previously identified as EPHB4 ligands (Noberini *et al*, 2012), we further investigated expression and function of EFNB2, given that EPHB4 likely interacts with EFNs expressed on adjacent cells in Caco-2 cell spheroids to fulfil its function. We found that EFNB2 is confined to the basolateral membrane below ZO-1 signals, thereby showing its apical exclusion (Figure 6B). This compartmentalized expression of EFNB2 is consistent with it being an EPHB4 ligand in Caco-2 spheroids. To determine if EFNB2 is required for Caco-2 spheroid morphogenesis, we depleted EFNB2 using two distinct shRNAs and found that it led to abnormal spheroid morphology in ∼70% of structures analyzed (p<0.0001; ANOVA with Tukey’s correction for multiple comparison) (Figure 6C-D), a proportion that is similar to what we found for aberrant structures formed by EPHB4-depleted Caco-2 cells (Figure 2). These observations are consistent with EFNB2/EPHB4 being a functional ligand-receptor pair in the regulation of Caco-2 cell spheroid morphology.

**Figure 6.**
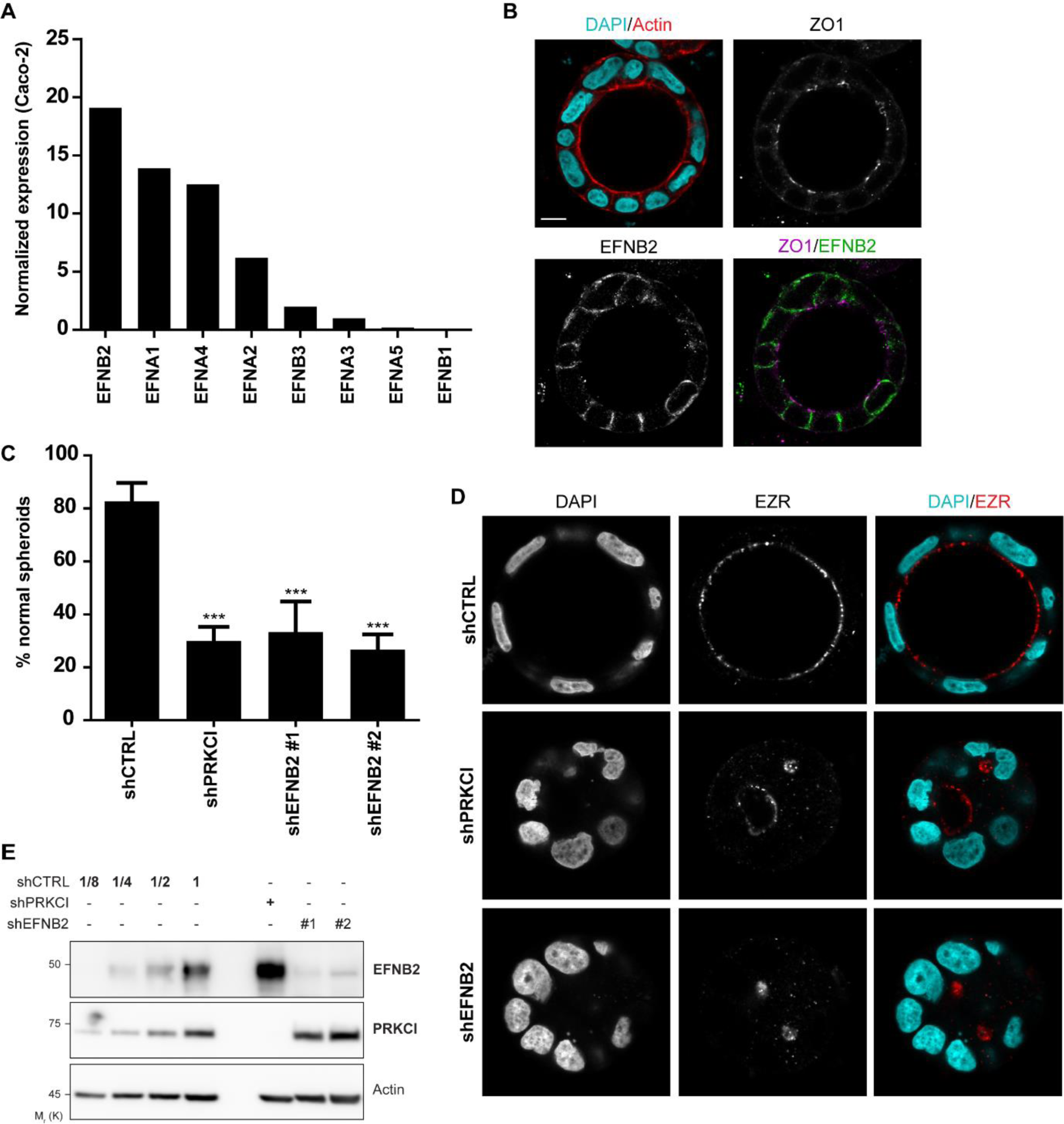
EFNB2 depletion leads to defects in Caco-2 spheroid formation. (A) RNA expression levels of all human EFNs in Caco-2 cells were extracted from the Human Protein Atlas database and displayed in a decreasing order. (B) Confocal images displaying a representative two examples of EFNB2 expression in 3D Caco-2 spheroid cysts. Single cells were seeded on Matrigel and grown for 6 days. Expression of apical (Actin) and tight junction (ZO-1) markers is shown, along with nuclear DAPI staining (scale bar: 15 µm). (C) Quantification of the percentage of Caco-2 spheroids displaying a wild-type morphology (i.e. single, round lumen judged by EZR staining, surrounded by a monolayer of cells) following depletion of EFNB2. The mean of three independent experimental replicates is shown. Error bars indicate SD (***p≤0,001; ANOVA with Tukey’s correction for multiple comparison; n=150). (D) Representative images of Caco-2 spheroids morphology following depletion of PKCι or EFNB2. EZR apical marker highlights the lumen border, along with nuclear DAPI staining. Single cells were seeded on Matrigel and grown for 6 days. (E) Western blot analysis of endogenous PKCι or EFNB2 following shRNA depletion. Two independent shRNAs were used against EPHRs. Successive dilutions of control are indicated to estimate depletion efficiency, where 1 is equivalent to 50 µg. Representative images of 3 experiments are shown.

## DISCUSSION

We report here that EPHA1 and EPHB4 receptors are expressed in both proliferative and differentiating Caco-2 cells. In polarized Caco-2 cell spheroids, EPHA1 and EPHB4 localize practically exclusively to the basolateral domain. Lateral localization of these receptors suggests a putative binding in *trans* to their membrane-tethered ephrin ligands also positioned at this subcellular region. Indeed, our data indicate that EFNB2 is not only enriched at the lateral membrane in Caco-2 cell spheroids, but is also essential for organizing proper spheroid morphology. Our observations are also consistent with previous reports of EPHA2 and EPHB4 being expressed at the basolateral membrane in MDCK cells and human pluripotent stem cell cysts, respectively (Harada *et al*, 2015; Wang *et al*, 2021). It remains a possibility that EPHRs at the basal domain associate in *cis* with ephrins, a mode of interaction that was previously detailed in different cellular contexts (Carvalho *et al*, 2006; Kao & Kania, 2011; Yin *et al*, 2004; Falivelli *et al*, 2013). Alternatively, EPHRs located at the basal membrane may function through crosstalk with other RTKs such as EGFR (Hanover *et al*, 2023; Hiramoto-Yamaki *et al*, 2010), or associate with matrix components, like fibronectin (Masuda *et al*, 2008).

We validated the protein levels of EPHA1, EPHA2, EPHA4 and EPHB4 in sub-confluent Caco-2 cells. The co-expression of several EPHR paralogs in the same tissue, yet the same cells is a frequent feature, and highlights EPHRs’ ability to play complementary or even partially redundant roles (Hafner *et al*, 2004). Interestingly, in differentiating Caco-2 cells, only EPHA1 and EPHB4 expression was maintained. Our observation that the depletion of either EPHA1 or EPHB4 is sufficient to impair spheroid morphogenesis supports the hypothesis that they have complementary, non-redundant roles. This is in accordance with our previous work, where we unraveled that dynamic protein complexes forming around EPHRs not only involve core (shared) signaling effectors but also components unique to specific EFN-EPHR pairs (Banerjee *et al*, 2022). This is further supported by our finding that EPHA1’s, but not EPHB4’s intracellular domain is required for spheroid morphogenesis and suggests that the two receptors act via distinct mechanisms.

We found through our rescue experiments that tyrosine kinase domain of EPHA1 and EPHB4 are not required for establishing and maintaining spheroid morphogenesis. This is hardly surprising, given that in some instances, EPHRs may execute non-catalytic functions (Liang *et al*, 2019). For example, EPHA4 kinase-dependent and kinase-independent functions are both involved in the formation of the anterior commissure in mouse, as kinase-dead EPHA4 rescued defects found in the *EPHA4* null animals (Kullander *et al*, 2001). Similar findings were reported for *EPHB2*^-/-^ animals, as a C-terminally truncated EPHB2 protein was able to rescue the synaptic plasticity and accompanying behavioral defects observed in the knockout animals (Grunwald *et al*, 2001). The distinction between kinase-dependent and kinase-independent functions for EPHB2 was also reported. While EPHB2 catalytic activity is required for cell proliferation in colon crypts in adult mice, it was dispensable for cell positioning/migration of Paneth cells relative to the crypt base (Genander *et al*, 2009). The latter required PI3K activation initiated by the receptor. In the case of EPHA2, ligand-independent activation of forward signaling was shown to rely on S897 phosphorylation by AKT (Miao *et al*, 2009), RSK (Zhou *et al*, 2015) or PKA (Barquilla *et al*, 2016); interestingly, the S897 phosphorylation was reported to coexist with kinase-dependent EPHA2 Tyr phosphorylation and canonical forward signaling (Barquilla *et al*, 2016). Finally, EPHA8 was reported to promote cell attachment to fibronectin independently of its catalytic activity, via the recruitment of PI3K to the plasma membrane (Gu & Park, 2001). These reports, combined with our present findings, imply a more intricate role for EPHRs beyond their traditional kinase-dependent signaling, highlighting their versatility in modulating cellular functions through diverse mechanisms. This nuanced perspective underscores the complexity of signal transduction networks and emphasizes the need to consider both catalytic and non-catalytic contributions of EPH receptors in deciphering their roles in various biological processes. This is further exemplified by the existence of two pseudo-kinases among the EPH family of receptors, namely EPHA10 and EPHB6 (Truitt & Freywald, 2011; Liang *et al*, 2021). Moreover, catalytically active EPHRs may display kinase-independent functions, exerting regulatory effects through scaffolding in forward signal transduction, or acting as ligands to initiate reverse signaling. Our data show that the entire intracellular domain of EPHB4, which also includes a SAM domain and a PBM, is dispensable. This suggests that EPHB4 may be strictly required for reverse signaling through one of its ephrin ligands. Accordingly, we found that EFNB2, the ligand displaying the best affinity for EPHB4 *in vitro* (Noberini *et al*, 2012), is required for spheroid morphogenesis. It is possible that A-type ephrin ligands EFNA1 and EFNA4, which are deemed as the most abundantly expressed in Caco-2 cells according to the Human Protein Atlas, are *bona fide* EPHA1 ligands *in vivo*, as they are *in vitro*(Noberini *et al*, 2012); this remains to be validated experimentally.

While our work did not pinpoint that the depletion of EPHA1 or EPHB4 led to major defects in the establishment or maintenance of cell polarity, we found that it led to a randomization of spindle orientation in developing spheroids. Numerous studies have shown that orientation of cell division contributes to maintain a single epithelial layer (Royer & Lu, 2011; Bergstralh & St Johnston, 2014). Intracellular factors aligning the mitotic spindle with the epithelial plane have been extensively described in the literature. However, very few membrane receptors were previously linked to mitotic spindle orientation. One example is the plexin-semaphorin combination, which regulates cell migration, proliferation, and differentiation in the nervous, immune, and bone systems (Worzfeld & Offermanns, 2014; Tamagnone, 2012). Plexins and semaphorins were identified as regulators of mitotic spindle alignment (Xia *et al*, 2015). Specifically, upon binding with semaphorins, plexins negatively regulate CDC42 activity and, consequently, mitotic spindle orientation. In the *Drosophila* optic lobe, EFN-EPHR signaling was shown to regulate mitotic spindle orientation (Franco & Carmena, 2019a). The authors demonstrated a requirement for parallel activation of both direct and reverse signaling in neuroepithelial cells of the *Drosophila* optic lobe. Null mutation of the unique *Drosophila Eph* leads to misoriented cell divisions and disrupts the localization and activation of aPKC, as well as the localization of key factors that are required for mitotic spindle anchoring, such as Dlg-1, Mud (NuMA in mammals), and Canoe (AFDN in mammals) (Franco & Carmena, 2019b, 2019a).

During cell division, certain proteins are relocated. An example is the protein Lgl in Drosophila, which is repositioned from the cortex to the cytoplasm (Bell *et al*, 2015; Carvalho *et al*, 2015; Moreira *et al*, 2019). The disruption of the basolateral distribution of RTK such as EGFR has been shown to result in missoriented mitotic spindle and tissue disorganization (Bañón-Rodríguez *et al*, 2014; Yoder *et al*, 1996). As our work suggests that EPH receptors may regulate mitotic spindle orientation, we validated the subcellular localization of EPH receptors during Caco-2 cell division. Our results showed that the localization of these two receptors is maintained at the membrane (Figure 3). Since their localization remains the same throughout the cell cycle, EPHA1 and EPHB4 could participate in the lateral anchoring of the mitotic spindle to the cell cortex during mitosis.

A number of studies have identified polarity proteins and intercellular junction proteins as interaction or proximity partners for EPHRs. Banerjee et al. characterized EPHA4, EPHB2, EPHB3 and EPHB4 receptor-dependent signaling networks in HEK293T cells using a BioID proximity labeling method and found the basolateral determinants SCRIB, DLG, LLGL1 and MARK2/3, the apical determinants PKCι, PARD6B and EPB41L5 and the junctional proteins AFDN, NECTIN2, CTNNA1, CTNND1 (p120) and OCLN as EPHRs partners (Banerjee *et al*, 2022). According to BioID results, most of these proteins associate with at least three EPHRs, suggesting that this association is conserved in the EPHR family. These data are supported by the observation that AFDN, SCRIB and MARK2 were identified as EPHA2 partners in epidermis cells (Perez White *et al*, 2017). EFNs were also found to bind polarity and junctional proteins; for example, OCLN and CLDN-4 were identified as EFNB1 partners (Fredriksson *et al*, 2015). In *Xenopus*, EFNB1 was reported to regulate tight junctions in epithelial cells, via its effect on the Par polarity complex (Lee *et al*, 2008). Our results suggest that these previously reported interactions do not influence epithelial cell polarization along the apical-basal axis *per se*, but rather impact on spindle orientation. Accordingly, many polarity proteins that are part of EPHRs’ protein networks were shown to control spindle orientation in epithelial cells (Jaffe *et al*, 2008; Qin *et al*, 2010; Rodriguez-Fraticelli *et al*, 2010b; Vorhagen & Niessen, 2014; Hao *et al*, 2010b; Gloerich *et al*, 2017; Hart *et al*, 2017; Porter *et al*, 2019). Altogether, our work sheds new light on the non-redundant, catalytically independent functions of EPHA1 and EPHB4 receptors in epithelial morphogenesis, which are likely to be of general biological relevance taken in the context of the previously published observations discussed above.

## ACKNOWLEDGMENTS

We thank C. St-Pierre for expert technical assistance with microscopy. This work was funded by grants from the Fonds de Recherche du Québec – Nature et Technologies (FRQNT) Team Grant program (254641 & 340951) to N.B., S.E. and P.L. N.L. held an Alexander-Graham-Bell Canada Graduate Scholarship, a FRQS Doctoral Award and a Frederick-Banting and Charles-Best Canada Graduate Scholarship from CIHR during the completion of this work. M.L. is supported by a salary award from the Fonds de Recherche du Québec-Santé (FRQS). N.B. holds a Canada Research Chair (Tier 2) in Cancer Proteomics.

## AUTHOR CONTRIBUTIONS

Investigation: N.L., A.S., F.J.-M.C., K.G., A.J.;

Resources: F.J.-M.C., S.L.B.;

Writing – Original Draft: N.L., P.L., N.B.;

Writing – Review & Editing: M.L., A.F., S.E., P.L., N.B.;

Visualization: N.L., F.J.-M.C.;

Supervision: M.C., M.L., A.F., S.E., P.L., N.B.;

Project Administration: P.L., N.B.;

Funding Acquisition: S.E., P.L., N.B.

## METHODS

### Cell culture, transfections and transductions

Caco-2 human colorectal adenocarcinoma cells and human embryonic kidney 293T cells (HEK293T) were cultured in Dulbecco’s Modified Eagle’s medium (DMEM, Thermo Fisher Scientific) high glucose supplemented with 10% fetal bovine serum (Sigma-Aldrich) at 37°C under 5% CO2.

Lentiviruses were produced using PEI co-transfection of 12 μg of pLKO constructs, 2 μg of pMD2.G (Addgene plasmid #12259, from Trono lab) and 6 μg of psPAX2 (Addgene plasmid #12260, from Trono lab). Culture media was changed 12h after transfection, then viral supernatants were collected 36h later and filtered through a 0.45 µm membrane. Cells were transduced for 16h prior to experiments and supplemented with 8 μg.ml^-1^ polybrene (Sigma-Aldrich).

Retrovirus for exogenous EPHRs expression in Caco-2 were produced as follows. Recombinant retroviruses were produced by transient transfection of 293GP21C cells (Ghani *et al*, 2007) with pMD2.G and MGF transfer vector using PEI. The culture supernatant was harvested on the second day after one medium change, spun down at 200 g for 5 min and filtered through a 0.45 µm membrane. Retroviruses were titrated with HT-1080 cells by flow cytometry (FACS). Caco-2 cells were transduced with a MOI of 1 for 72h prior to experiments and supplemented with 8 μg.ml^-1^ polybrene (Sigma-Aldrich).

### Constructs and sequences

ShRNA sequence targeting human PKCι (TRCN0000219727), EPHA1 (TRCN0000314913 and TRCN0000314914), EPHB4 (TRCN0000314827 and TRCN0000001773) and EFNB2 (TRCN0000058424 and TRCN0000285596) were inserted downstream of the U6 promoter in the pLKO vector (Addgene plasmid #8453, from Trono lab). Control vector was pLKO with a non-targeting shRNA sequence (5’-CCG GTC CTA AGG TTA AGT CGC CCT CGC TCG AGC GAG GGC GAC TTA ACC TTA GGT TTT T-3’). EGFP sequence was inserted in all pLKO vectors to detect infected cells.

Mutants of EPHA1 and EPHB4 were generated using the Q5-site directed mutagenesis kit (New England BioLabs): EPHA1-KD (K656A), EPHB4-KD (K647A), EPHA1-mLBD-GFP (EPHA1/R104E), EPHB4-mLBD-GFP (EPHB4/L95R). EPHA1-ΔSAM/PBM (1-910), EPHA1-ΔICD (1-592), EPHB4-ΔSAM/PBM (1-904) and EPHB4-ΔICD (1-583).

The retroviral MFG plasmid backbone used in this study was derived from Luc1 (Qiao *et al*, 2002) and contained an encephalomyocarditis virus internal ribosomal entry site sequence linked to a Tomato sequence instead of the luciferase gene. EPHR sequences with GFP-tag in C-ter were inserted with the NEBuilder HiFi DNA Assembly Cloning Kit (New England BioLabs) upstream of the IRES. Both genes are thus under the control of the Moloney murine leukemia virus (MoMLV) long terminal repeat (LTR) sequences.

Human EPHA1 and EPHB4 were made resistant to the shRNA by Q5-site directed mutagenesis kit in the MFG vector. The WT EPHA1 sequence GT**C** AA**T** GG**C** CT**T** GA**A** CCT was modified to GT**<u>G</u>** AA**<u>C</u>** GG**<u>A</u>** CT**<u>G</u>**GA**<u>G</u>** CCT for TRCN0000314914 and the WT EPHB4 sequence AA**T** GG**G** AG**A** GA**A** GC**A** GAA was modified to AA**<u>C</u>** GG**<u>C</u>**AG**<u>G</u>** GA**<u>G</u>** GC**<u>T</u>** GAA for TRCN0000001773.

### Cell lysis and affinity purification

Cells were washed once with ice-cold PBS. Caco-2 cells were lysed in KLB buffer (20 mM Tris-HCl pH 7.4, 150 mM NaCl, 1 mM EDTA, 0.5% sodium deoxycholate) supplemented with protease inhibitors (P8340, 1:100, Sigma). Caco-2 cells were incubated with lysis buffer in the Petri dish at 4°C for 15 min, then scraped before a 20 min centrifugation at 20 000 g. HEK293T cells were scraped, then incubated on ice for 15 min. Protein concentrations were measured and normalized using a BCA assay (Thermo Fisher Scientific).

Lysates were incubated with GFP-Trap agarose beads (Chromotek) for 2h at 4°C with rotation. Phosphatase inhibitors (Cocktail 2, 1:100, Sigma-Aldrich) and protease inhibitors (P8340, 1:100, Sigma) were added in lysis buffer. Beads were washed three times with lysis buffer then boiled in Laemmli buffer (Laemmli, 1970).

### Western blotting

Lysates in Laemmli buffer were loaded on SDS-PAGE gels for protein separation and nitrocellulose membranes were used for transfer. Loading was validated by Ponceau S (Sigma-Aldrich). Primary antibodies used were: goat anti-EPHA1 (R&D, 1:200, #AF638), goat anti-EPHB4 (R&D, 1:200, #AF3038), mouse anti-EPHA4 (BD Biosciences, 1:1000, #610471), mouse anti-EPHA2 (Millipore-Sigma, 1:500, #05-480), mouse anti-Actin (Cell Signaling Technology, 1:1000 #3700), mouse anti-PKCι (BD, 1:500, #610175), rabbit anti-GFP (Thermo Fisher Scientific, 1:1000 #A-11122) and rabbit anti-EFNB2 (R&D, 0.2 µg/µL, #NBP1-84830). Secondary antibodies used were: goat anti-rabbit IgG HRP (Cell Signaling Technology, 1:10 000, #7074), rabbit anti-goat IgG HRP (Thermo Fisher Scientific, 1:10 000, #31402) and horse anti-mouse IgG HRP (Cell Signaling Technology, 1:10 000, #7076). Signal was revealed using Clarity western ECL substrate or Clarity Max Western ECL Substrate (Bio-Rad) and acquired on an Amersham Biosciences Imager 600RGB (Cytiva).

### Caco-2 spheroids formation

To produce Caco-2 cysts, single cells were plated on top of a 20 μL pre-coat of 80% Matrigel Growth Factor Reduced (Fisher Scientific, CB-40230A) and 20% Collagen I, Bovine (Fisher, CB-4023) in a µ-Slide 8 Well Ibidi plate (Ibidi, 80826). Trypsinized cells were resuspended in culture medium supplemented with 2% Matrigel Growth Factor Reduced; 10 000 cells were seeded per well. Cells were grown for two or six days before fixation for immunofluorescence. For loss-of-function and overexpression experiments, cells were infected 24 h prior to seeding on Matrigel. For rescue experiments, cells were infected with retroviruses 48 h before lentivirus infection. For Western blot analyses, 2D cultures were grown in parallel for two or six days. For the two-day 3D cultures, 400 000 cells were seeded in 6-well plates. For the six-day 3D cultures, 400 000 cells infected with shEPHB4 and shCTRL as well as 550 000 cells infected with shEPHA1 were seeded in 10 cm Petri dishes.

### Immunofluorescence

For Caco-2 spheroids, cells were fixed after two or six days of culture on Matrigel with 4 % paraformaldehyde for 15 to 60 min at room temperature or at 37°C. 15 min fixation was for EPHB4 immunofluorescence only. Cells were washed 3 times for 15 min with PBS, then permeabilized and blocked for 1 h at room temperature in 0.5% Triton X-100 (Sigma-Aldrich), 10% donkey serum (Sigma) in PBS. Cells were incubated with goat anti-EPHA1 (R&D, 1:200, #AF638), goat anti-EPHB4 (R&D, 1:200, #AF3038), rabbit anti-Ezrin (Cell signaling, 1:200, #3145S), mouse anti-ZO-1 (Thermo Fisher Scientific, 1:50, #33-9100), mouse anti-Par6 (Santa Cruz, 1:100, #166405), mouse anti-E-cadherin (BD Biosciences, 1:200, #610181), rabbit anti-E-cadherin (Cell Signaling, 1:200, #3195), goat anti-Scrib (Santa Cruz, 1:200, #11049), rabbit anti-GFP (Thermo Fisher Scientific, 1:100, #A-11122), rabbit anti-α-tubulin (Abcam, 1:200,

#18251) and rabbit anti-EFNB2 (R&D, 3:200, #NBP184830) overnight at 4°C. Following 3 washes of 15 min in blocking/permeabilizing solution, cells were incubated for 2 h with Alexa Fluor 488 donkey anti-goat IgG (Invitrogen, 1:400, #A11055), Alexa Fluor 568 donkey anti-mouse IgG (Invitrogen, 1:400, #A10037), Alexa Fluor 647 donkey anti-rabbit IgG (Invitrogen, 1:400, #A31573), Alexa Fluor 488 donkey anti-rabbit IgG (Thermo Fisher Scientific, 1:400, #A21206), Alexa Fluor 647 donkey anti-mouse IgG (Cedarlane, 1:400, 715-605-150), Alexa Fluor 568 donkey anti-rabbit IgG (Invitrogen, 1:400, #A10042), Alexa Fluor 647 donkey anti-goat IgG (Thermo Fisher Scientific, 1:400, #A21447) and/or DAPI (1:5000, Sigma-Aldrich). Cells were washed 3 times for 15 min in PBS and left in PBS until image acquisition. Images were acquired on a confocal laser scanning microscope (Olympus FV1000 or Zeiss LSM 900) using PlanApo 40x (numerical aperture 0.90) objectives or C-Apochromat 40x 1.1 NA Water. Images were processed with ImageJ. Division angle measurements were taken using the angle tool in ImageJ.

### Statistics

GraphPad Prism 8 was used for statistical analysis. Statistical differences between conditions for loss-of-function and overexpression experiments were assessed with one-way ANOVA followed with Tukey’s correction for multiple comparison. Statistical differences between conditions for rescue experiments were assessed with one-way ANOVA with Dunnett’s correction for multiple comparison and each condition was compared to a single control, as opposed to all datasets. Statistical differences in the angle of mitotic spindle were assessed by a Kruskall-Wallis test (all compared to control) as the samples were small.

## SUPPLEMENTAL FIGURES

**Figure S1.**
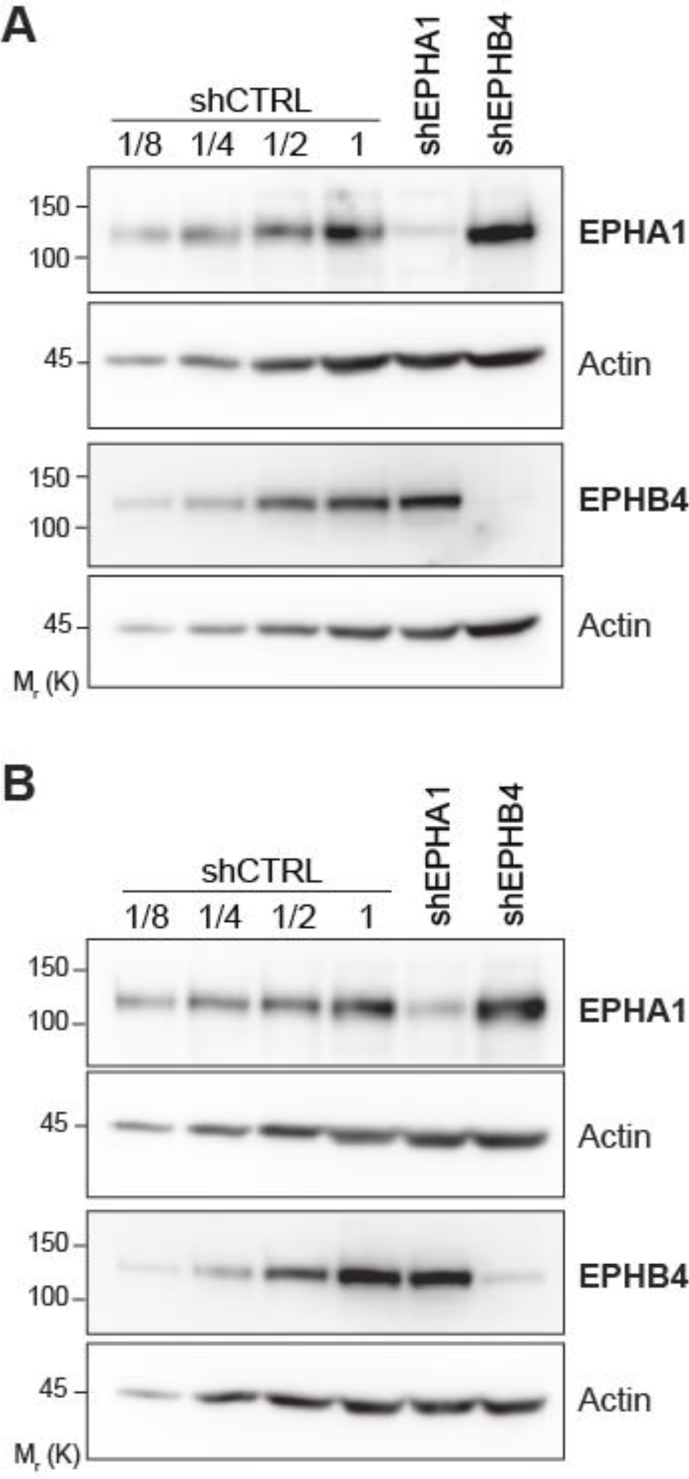
Western blot analyses complementary to Figures 3 and 4. Western blot analysis of endogenous EPHA1 and EPHB4 following shRNA depletion in Caco-2 cells cultured as monolayers. Successive dilutions of control are indicated to estimate depletion efficiency, where 1 is equivalent to 50 µg. Representative images of at least 3 experiments are shown, (A) corresponding to Figure 3 and (B) corresponding to Figure 4.

**Figure S2.**
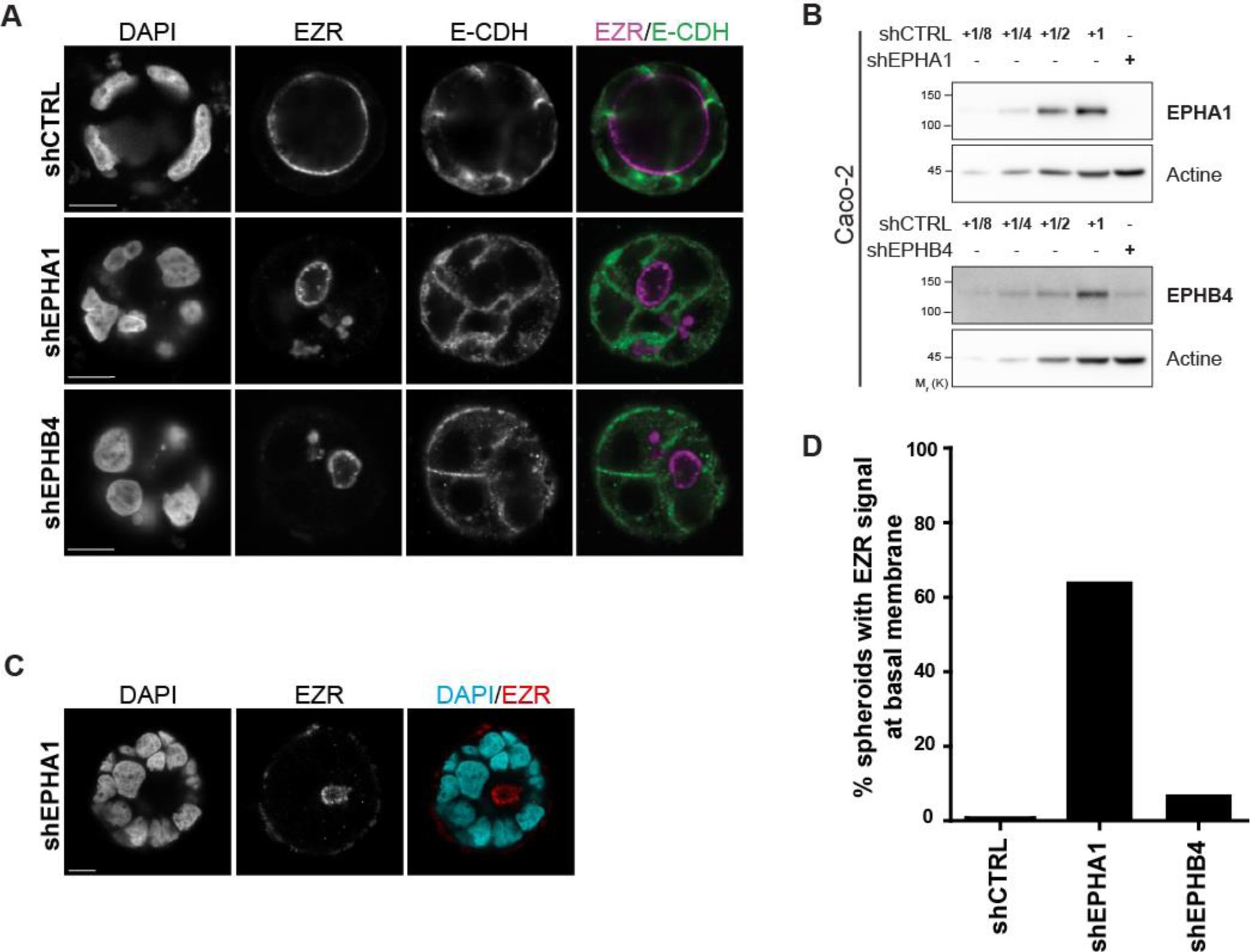
Depletion of EPHA1 leads to delocalization of EZR to the basal membrane of Caco-2 spheroids. (A) Representative images of Caco-2 spheroids showing EZR subcellular localization following EPHA1 or EPHB4 depletion. Single cells were seeded on Matrigel and grown for 6 days. E-CDH highlights basolateral membranes, along with nuclear DAPI staining (scale bar: 12 µm). (B) Western blot analysis of endogenous EPHA1 and EPHB4 in Caco-2 cells cultured as monolayers. Successive dilutions of control are indicated to estimate depletion efficiency, where 1 is equivalent to 50 µg. Representative images of at least 3 experiments are shown. (C) Representative images of Caco-2 spheroids showing EZR at the basal membrane after EPHA1 depletion. Single cells were seeded on Matrigel and grown for 6 days (scale bar: 10 µm). (D) Quantification of the percentage of spheroids with EZR signal at the basal membrane (n=200 for shCTRL, n=650 for shEPHA1 and n=500 for shEPHB4).

**Figure S3.**
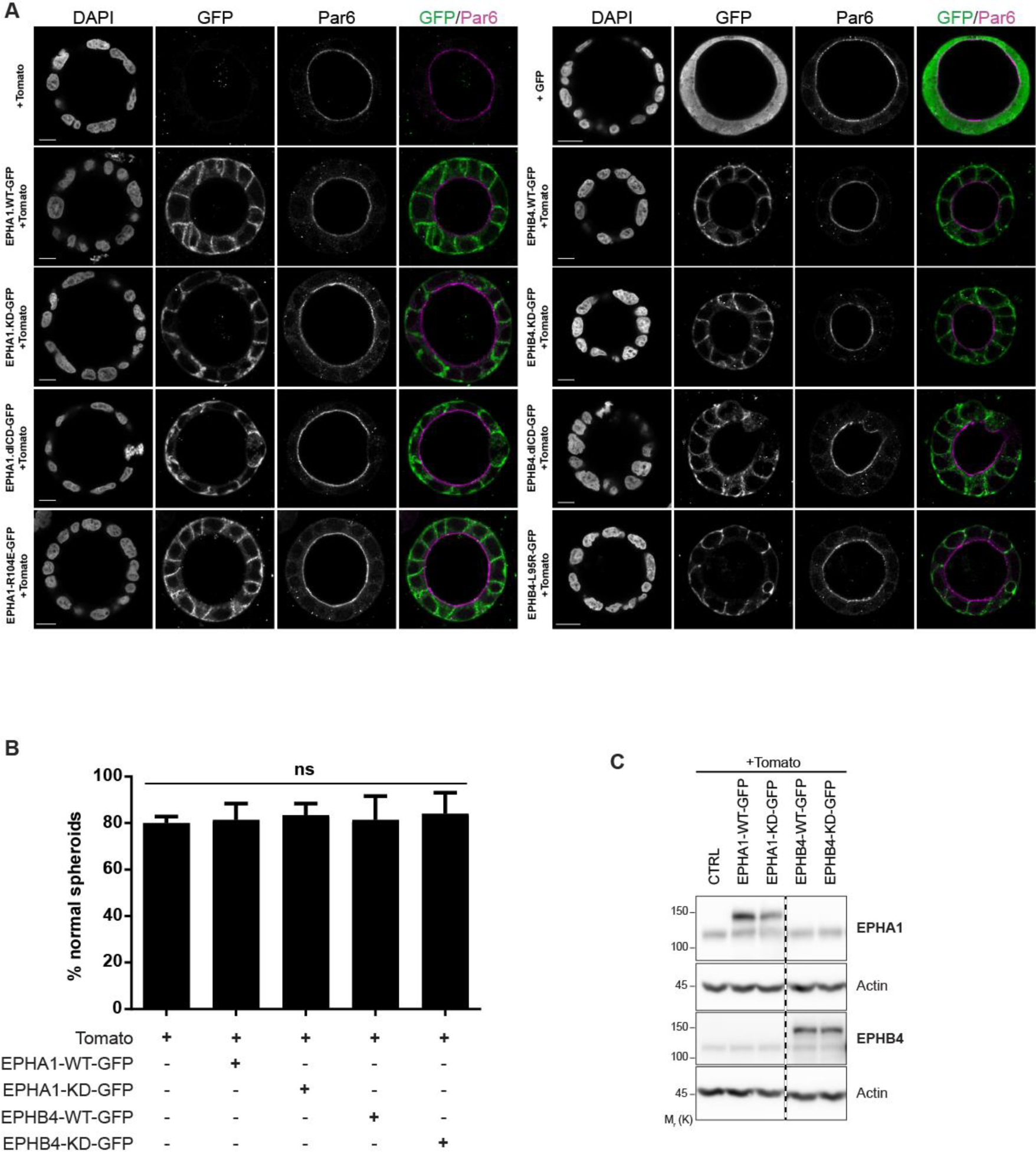
Overexpression of EPHA1 or EPHB4 does not affect spheroid morphogenesis. (A) Representative images of Caco-2 spheroids following EPHA1 or EPHB4 exogenous expression (GFP). PARD6 (Par6) apical marker highlights the lumen border, along with nuclear DAPI staining. Single cells were seeded on Matrigel and grown for 6 days (scale bar: 15 µm). (B) Quantification of the percentage of Caco-2 spheroids displaying a wild-type morphology (i.e. single, round lumen judged by EZR staining, surrounded by a monolayer of cells) following EPHA1/B4-WT-GFP and EPHA1/B4-KD-GFP overexpression. The mean of 3 independent experimental replicates is shown. Error bars indicate SD (***p≤0,001; ANOVA with Tukey’s correction for multiple comparison; n=400 for control and n=150 for each EPHR-WT/KD-GFP). (C) Western blot analysis of endogenous and exogenous (WT and KD, GFP tagged) EPHA1 and EPHB4 expression. Representative images of 3 experiments are shown.

